# Unraveling Vulnerabilities in Endocrine Therapy-Resistant HER2+/ER+ Breast Cancer

**DOI:** 10.1101/2023.08.21.554116

**Authors:** Shaymaa Bahnassy, Hillary Stires, Lu Jin, Stanley Tam, Dua Mobin, Manasi Balachandran, Mircea Podar, Matthew D. McCoy, Robert A. Beckman, Rebecca B. Riggins

## Abstract

**Background:** Breast tumors overexpressing human epidermal growth factor receptor (HER2) confer intrinsic resistance to endocrine therapy (ET), and patients with HER2/ estrogen receptor-positive (HER2+/HR+) breast cancer (BCa) are less responsive to ET than HER2−/ER+. However, real-world evidence reveals that a large subset of HER2+/ER+ patients receive ET as monotherapy, positioning this treatment pattern as a clinical challenge. In the present study, we developed and characterized two distinct *in vitro* models of ET-resistant (ETR) HER2+/ER+ BCa to identify possible therapeutic vulnerabilities.

**Methods:** To mimic ETR to aromatase inhibitors (AI), we developed two long-term estrogen-deprived (LTED) cell lines from BT-474 (BT474) and MDA-MB-361 (MM361). Growth assays, PAM50 molecular subtyping, genomic and transcriptomic analyses, followed by validation and functional studies, were used to identify targetable differences between ET-responsive parental and ETR-LTED HER2+/ER+ cells.

**Results:** Compared to their parental cells, MM361 LTEDs grew faster, lost ER, and increased HER2 expression, whereas BT474 LTEDs grew slower and maintained ER and HER2 expression. Both LTED variants had reduced responsiveness to fulvestrant. Whole-genome sequencing of the more aggressive MM361 LTED model system identified exonic mutations in genes encoding transcription factors and chromatin modifiers. Single-cell RNA sequencing demonstrated a shift towards non-luminal phenotypes, and revealed metabolic remodeling of MM361 LTEDs, with upregulated lipid metabolism and antioxidant genes associated with ferroptosis, including *GPX4*. Combining the GPX4 inhibitor RSL3 with anti-HER2 agents induced significant cell death in both the MM361 and BT474 LTEDs.

**Conclusions:** The BT474 and MM361 AI-resistant models capture distinct phenotypes of HER2+/ER+ BCa and identify altered lipid metabolism and ferroptosis remodeling as vulnerabilities of this type of ETR BCa.

## INTRODUCTION

Globally, breast cancer (BCa) remains the leading cause of cancer-related mortality in women (1). Human epidermal growth factor receptor positive/ hormone receptor positive (HER2+/HR+) tumors account for approximately 10% of all BCa cases and 60% of HER2+ tumors (2). HER2+/ estrogen receptor positive (ER+) and HER2+/ER-BCa have distinct biology and are proposed as two distinct disease subtypes (3,4). Prior and recent studies reported favorable prognosis and overall survival (OS) as well as lower-grade tumors in HER2+/ER+ BCa compared to HER2+/ER-BCa patients (5–8). However, HER2+/ER+ BCa has low response rates to anti-HER2 therapy compared to HER2+/ER-tumors (9,10), have a limited antiproliferative response to endocrine therapy (ET), and are at a higher risk of recurrence (2,10,11).

Concurrent blockade of both HER2 and ER pathways in patients with advanced/metastatic HER2+/ER+ BCa is becoming increasingly evident as an effective treatment strategy. In clinical trials, postmenopausal women receiving combinations of ET plus anti-HER2 as first-line therapy showed improvements in progression-free survival (PFS) and/or other clinical benefits (12–14). Likewise, a retrospective study of the national cancer database showed the highest 5-year OS among those receiving this combination therapy vs. ET alone, chemotherapy alone or anti-HER2 + chemotherapy (15). Unfortunately, outside of clinical trials a large subset (36.7-60%) of HER2+/ER+ patients receive hormonal treatment as a monotherapy (15,16). Although current guidelines recommend anti-HER2 agents (trastuzumab + pertuzumab; TP) plus chemotherapy as first-line standard of care for advanced HER2+ disease irrespective of HR status, endocrine monotherapy remains an option for selected HER2+/ER+ patients that have low disease burden or intolerance to chemotherapy (17,18). Moreover, antibody-based anti-HER2 therapy is mainly given to patients with primary HER2+/ER+ BCa for one year, while ET is usually given longer (at least five years) (2). Thus, residual HER2+/ER+ disease after surgery, not subjected to anti-HER2 therapy, remains at high risk of incomplete response or resistance to prolonged ET.

A shorter time to recurrence in patients with HER2+ disease, compared to HER2-, treated with single-agent ET as tamoxifen or anastrozole was reported (19). Likewise, compelling evidence from preclinical and clinical data strongly suggests that HER2 overexpression confers intrinsic resistance to ET and that HER2+/ER+ BCa are less responsive to ET than HER2−/ER+ tumors (2,10,20). Multiple studies have further established that bi-directional crosstalk between ER and HER2 signaling pathways mediates resistance to ET (21–23). While acquired ET resistance is extensively studied in HER2−/ER+ BCa, preclinical models of HER2+/ER+ BCa that have acquired resistance to ET are lacking. Existing models of resistance have been generated by genetic manipulations of ER and/or HER2, for example (24,25).

Collectively, there is an unmet need to develop, characterize and study therapeutic responses of ET-resistant (ETR) HER2+/ER+ models to HER2- and other ER-directed therapies. In this study, we establish ETR variants through long-term estrogen deprivation (LTED) to mimic resistance to aromatase inhibitors (AI) from two HER2+/ER+ BCa cell lines (BT474 and MDA-MB-361, (26,27)). To identify therapeutic vulnerabilities between AI-responsive-parentals and ETR-LTEDs of HER2+/ER+ BCa cell lines, we performed growth assays, PAM50 molecular subtyping, genomic and transcriptomic analyses. Our data shows that the BT474 and MM361 ETR models capture distinct phenotypes of HER2+/ER+ BCa and highlight altered lipid metabolism and ferroptosis remodeling as features of this ETR BCa.

## METHODS

### Treatments, Cell Lines, and Cell Culture

Trastuzumab and pertuzumab were obtained from Genentech, SYTOX green (S7020) and Hoechst 33342 (H1399) from ThermoFisher, fulvestrant (S1191) from Selleck Chemicals, β-estradiol (E8875) from Sigma Aldrich and RSL3 (HY-100218A) from MedChemExpress.

MDA-MB-361 (MM361; RRID:CVCL_0620) and BT-474 (BT474; RRID:CVCL_0179) human breast carcinoma cell lines were obtained from the Tissue Culture and Biobanking Shared Resource at Lombardi Comprehensive Cancer Center and were routinely checked for *Mycoplasma* contamination. Both cell lines were cultured in improved Minimum Essential Medium (IMEM; Gibco) media supplemented with 10% FBS at 37°C in a humidified atmosphere containing 5% CO_2_. We generated two LTED variants A and B cells from their corresponding parental cell lines by chronically passaging parental cell lines in phenol red-free IMEM media (Gibco) supplemented with 10% charcoal-stripped bovine serum (CSS; Vita Scientific) for over six months. After developing resistance, the derived cells were used and continuously cultivated in 10% CSS phenol red-free IMEM.

### Growth Assays

Differences in growth kinetics between parental and corresponding LTEDs were evaluated by the trypan blue exclusion assay. Cells were seeded at a density of 100,000 cells/well in 24-well plates. After trypsinization and staining with trypan blue, live cell counts using the Countess II Automated Cell Counter (ThermoFisher Scientific) were recorded on days 0 (24 h), 2, 4, 6, 8, and 10 from cell seed.

For crystal violet assays, parental and LTED cells were seeded in 96-well plates at 5,000 cells/well. Forty-eight hours later, cells were subjected to indicated treatments for six additional days. Treatments were replenished three days after treatment onset. At the end of the experiments, cells were stained with 0.5% crystal violet in 25% methanol. Once plates were dried, the stain was resolubilized with citrate buffer, and absorbance measurements were obtained from the ELx808 plate reader (BioTek).

### Cell Death Quantification

Cells were seeded in 24-well plates at a cell density of 150,000 cells/well, incubated overnight, then subjected to treatments as indicated. After 72 h of treatments, cells were stained with both Hoechst 33342 (0.1 μg/mL) to monitor total cell number, and Sytox green (5 μM) to monitor dead cells. Images were acquired using the Olympus IX71 microscope and subsequently analyzed by Image J. Percentage cell death was calculated as Sytox green cell number over total cell number.

### Western Blot Analysis

Whole-cell protein extracts were denatured, resolved on NuPAGE 4-12% Bis-Tris gels (ThermoFisher), and either transferred onto nitrocellulose membranes using iBlot 2 dry transfer apparatus (ThermoFisher) or PVDF membranes using the BioRad wet transfer apparatus. After blocking, blots were probed overnight, and the following primary antibodies were used: ER (Cell Signaling Technology Cat# 8644, RRID:AB_2617128), EGFR (Cell Signaling Technology Cat# 2232, RRID:AB_331707), HER2 (Cell Signaling Technology Cat# 2242, RRID:AB_331015), pHER2 (Cell Signaling Technology Cat# 2241, RRID:AB_2099407), HER3 (Cell Signaling Technology Cat# 12708, RRID:AB_2721919), HER4 (Cell Signaling Technology Cat# 4795, RRID:AB_2099883), AKT (Cell Signaling Technology Cat# 4691, RRID:AB_915783), pAKT (Cell Signaling Technology Cat# 9271, RRID:AB_329825), β-actin (Cell Signaling Technology Cat# 3700, RRID:AB_2242334), vinculin (Cell Signaling Technology Cat# 13901, RRID:AB_2728768), 4-HNE (Abcam Cat # ab46545, RRID: AB_722490), GPX4 (Abcam Cat# ab125066, RRID:AB_10973901), MDA (Thermo Fisher Scientific Cat# MA5-27559, RRID:AB_2735264) and GAPDH (Proteintech Cat# 60004-1-Ig, RRID:AB_2107436). After one hour of incubation with secondary antibody (either anti-mouse (Cell Signaling Technology Cat# 7076, RRID:AB_330924) or -rabbit (Cell Signaling Technology Cat# 7074, RRID:AB_2099233)) and washes, antigen-antibody complexes were detected by the chemiluminescence WesternBright ECL Detection Reagent (Advansta) and imaged using the Amersham Imager 600 (GE Healthcare Life Sciences).

### Real-time PCR (qRT-PCR)

Total RNA of biological triplicates was extracted from cells using the PureLink RNA Mini Kit (ThermoFisher) and converted to cDNA with iScript cDNA Synthesis Kit (BioRad) according to manufacturer’s instructions. Expression of target genes with specific primers (sequences listed in Table S1 (28); synthesized by IDT) was measured by RT-qPCR using the iTaq Universal SYBR Green Supermix (BioRad) and QuantStudio 12K Flex Real-Time PCR System (ThermoFisher). Data were normalized to the ß-actin housekeeping gene and analyzed by the ΔΔCT method.

### Single-Cell RNA Sequencing (scRNAseq)

The scRNAseq was performed using the Drop-seq approach described by Macosko *et al*. (29) with updates and detailed workflow from the McCarroll lab at Harvard Medical School (https://mccarrolllab.org/dropseq/). To generate the droplet emulsion and cell encapsulation we used a Dolomite-Bio μEncapsulation system (Dolomite-Bio) according to manufacturer’s instructions. Barcoded beads were purchased from Chemgenes, as specified by (29). Briefly, trypsinized MM361 parental and LTED cells were washed, and resuspended in PBS-0.01% BSA. The single-cell suspension, bead suspension, and the droplet generation oil were loaded into their respective containers and connected via high precision pumps to the scRNA chip as part of the μEncapsulation system. The flow rates were adjusted so that each droplet generated encapsulated one bead and one cell and monitored under the system’s high speed digital microscope. Droplets were collected, and a small aliquot was examined microscopically to ensure uniformity of bead size and occupancy. The beads were washed and used for reverse transcription/cDNA synthesis and PCR according to the drop-seq workflow. The PCR products were purified, pooled, and quantified in a BioAnalyzer High Sensitivity Chip (Agilent). Sequencing libraries were prepared using the Nextera XT DNA sample prep kit (Illumina Inc) following manufacturer’s protocol. The libraries were purified, quantified, and sequenced by GenWiz on an Illumina High-Seq instrument using 2×150 nt reads.

Raw data was imported into the Seurat R package (30). All cells with unique feature counts between 200 and 2500 and a percentage of mitochondrial reads less than 5% were selected for further analysis. After scaling, dimensional reduction, and cell type specific marker identification with default parameters, we discovered 7 different clusters among LTED and parental cell lines. Differentially expressed genes (p < 0.05) were calculated between all three experimental groups. A combined LTED and parental cell lines UMAP was generated with 10 PC and 0.5 resolution. We also performed PAM50 molecular classification using genefu R package (31) at single-cell resolution; a treatment split UMAP was generated with the same settings.

### Gene Set Enrichment Assay (GSEA)

Differentially expressed genes were matched against the REACTOME signature from the Human Molecular Signature Database (MSigDB) using the GSEA portal (http://software.broadinstitute.org/gsea/index.jsp) (32), with false discovery rate (FDR) q-values <0.05. Top 20 enriched pathways and their corresponding - log (p-value) were graphed.

### Whole Genome Sequencing (WGS)

DNA was extracted from cells using the DNeasy Blood and Tissue Kit (Qiagen) according to manufacturer’s instructions. The Genomics and Epigenomics Shared Resource (GESR) at Georgetown University Medical Center performed the WGS. Paired-end, indexed libraries for human WGS were constructed from 1.0 μg of gDNA using the TruSeq DNA PCR-Free Library Prep Kit (Illumina) according to the manufacturer’s instructions. Briefly, DNA was fragmented using a Covaris M220 Focused-ultrasonicator (Covaris) using settings for a 350-bp insert size. Library quality was assessed with a BioAnalyzer 2100 using the High Sensitivity DNA kit (Agilent Technologies). The libraries were quantified using the Kapa Library Quantification Kit Illumina Platforms (Kapa Biosystems). The denatured and diluted libraries were sequenced on a NextSeq 550 System (Illumina) using v2.5 High Output 300 cycle kit with 1% PhiX to an average sequencing depth of 50x coverage.

The quality of raw sequence data (fastq or fasq.gz files) was checked by FastQC v0.11.9 (33), and Cutadapt v3.5 was used for adapter trimming of raw data (34). After trimming, reads with low quality (quality score < 33, error rate > 10%) and lengths less than 25 bp were eliminated. Processed reads were then aligned to GRCh38 reference sequence using the bwa v0.7.17 paired-end mode (35). Mutation detection was conducted in Genome Analysis Toolkit (GATK) v4.1.9.0 (36) following the best practice for variant calling workflow. SigProfiler (37) on COSMIC Mutational Signatures version 3.2 was used to classify single base substitution (SBS) in WGS data.

### Statistical Analysis

All results are expressed as mean ± SEM. Prism 9 (GraphPad Software) was used for data analysis. Unless otherwise noted, one-way ANOVA followed by Tukey’s or Dunnett’s multiple comparison tests were employed to evaluate statistical significance between groups and *p*-values < 0.05 were considered statistically significant.

## RESULTS

### Patients of the HER2+ BCa subtype expressing high levels of ESR1 exhibit lower risk for metastasis

We analyzed a publicly available database to evaluate patients’ survival rates of HER2+ BCa stratified by their HR status. Regardless of HR status, five-year survival rates from the SEER database for patients with HER2+ BCa decrease with disease progression and are the lowest for late (distant)-stage HER2+ BCa (**Fig. 1A**). Next, we specifically focused on ER and compared survival rates of HER2+ BCa patients as stratified by high and low estrogen receptor alpha gene (*ESR1*) expression. Ten-year survival data from the Kaplan-Meier Plotter database (38) show no difference in overall survival (OS) and relapse-free survival (RFS) between patients expressing high or low levels of *ESR1* (**Fig. 1B** and **1C**, respectively). On the contrary, patients expressing high levels of *ESR1* show higher probability for distant metastasis-free survival (DMFS) (logrank *P* = 0.035, **Fig. 1D**), and thus lower risk for metastasis. This lower susceptibility for metastasis suggests correlation of high ER expression with better prognosis within the advanced setting of HER2+ BCa disease.

**Figure 1.**
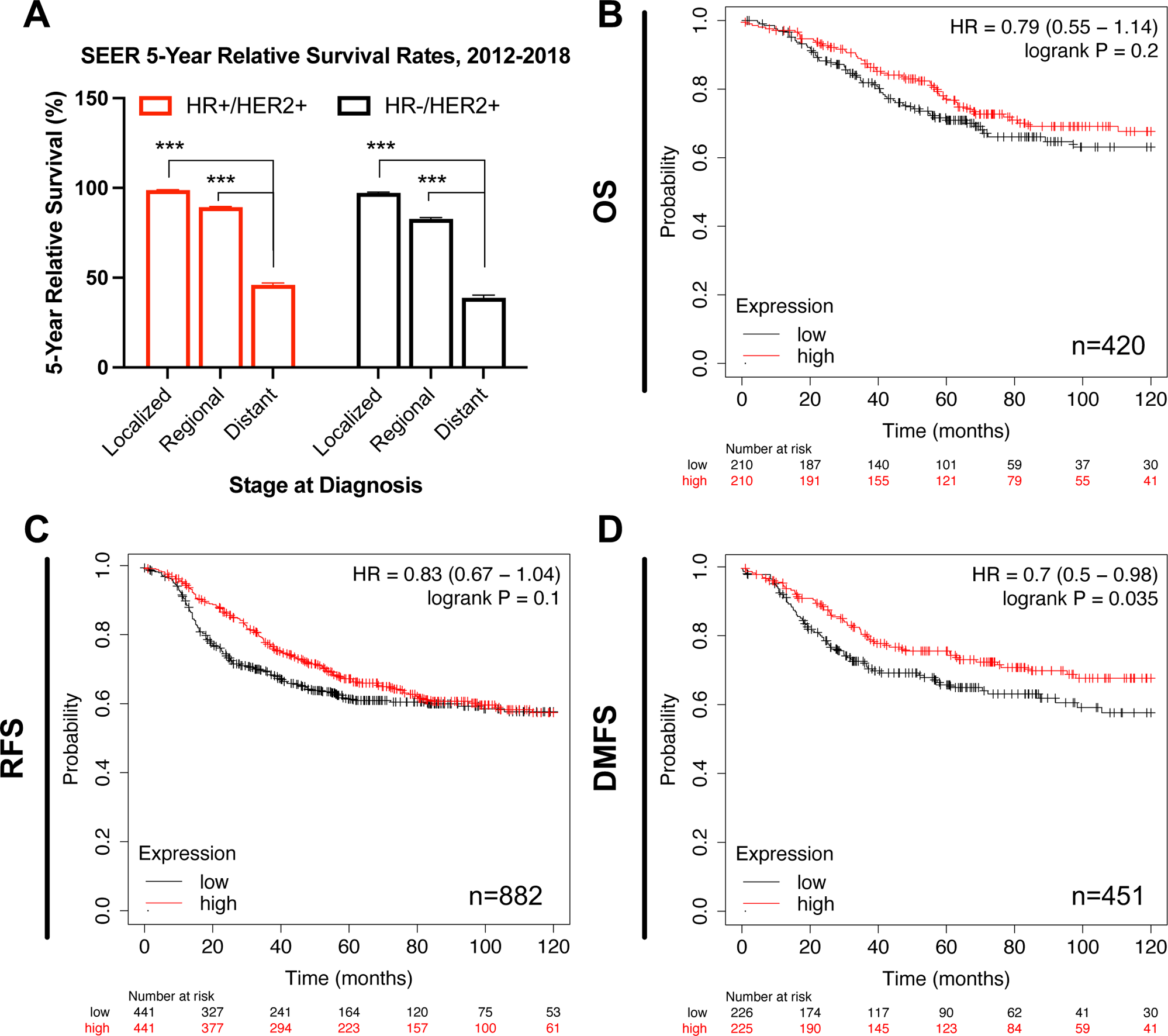
Survival analysis for HER2+ BCa regarding their ER status. **(A)** Compared to early-stage disease, five-year survival rates from the SEER database (https://seer.cancer.gov/statistics-network/explorer, accessed on 6 February 2023) are lowest for patients with late (distant)-stage HER2+ BCa subtype, regardless of their HR status. Error bars represent SEM. One-way ANOVA followed by Tukey’s multiple comparison was performed to compare between groups; \*\*\**p*L<L0.001 denotes statistically significant. **(B-D)** Patients of the HER2+ BCa subtype expressing high levels of *ESR1* have a significantly lower probability for metastasis. Ten-year survival curves for HER2+ BCa patients stratified by high and low gene expression of *ESR1* (205225_at). Plots were generated from KM Plotter database for BCa (www.kmplot.com, accessed on 2 February 2023) and show hazard ratio (HR) at 95% confidence, log-rank P values and number of patients (n). OS: Overall survival; RFS: Relapse-free survival; DMFS: Distant metastasis-free survival.

### Growth patterns and responses to HER2- and ER-targeted therapies of HER2+/ER+ LTEDs differ from their parental counterparts

To mimic acquired ETR to AI, we generated two LTED variants (A and B) from each of the two HER2-amplified, HER2+/ER+ BCa cell lines (**Fig. 2A** and **2B**): BT474 and MM361 (26,27). We first compared proliferation behaviors of LTEDs versus their corresponding parental cell lines grown in media supplemented with either FBS or CSS (CSS represents short-term hormone starvation). BT474 LTEDs had significantly lower growth rates than their respective parentals grown in FBS media but were not different from parentals in CSS media (**Fig. 2C** left panel). On the other hand, MM361 LTEDs grew faster than parentals in CSS and FBS media (**Fig. 2C** right panel). Increased growth rates of MM361 LTEDs in CSS media are suggestive of hormone-independent growth and a more aggressive nature (LTEDs being more aggressive than parentals), consistent with the isolation of MM361 from brain metastatic BCa (39).

**Figure 2.**
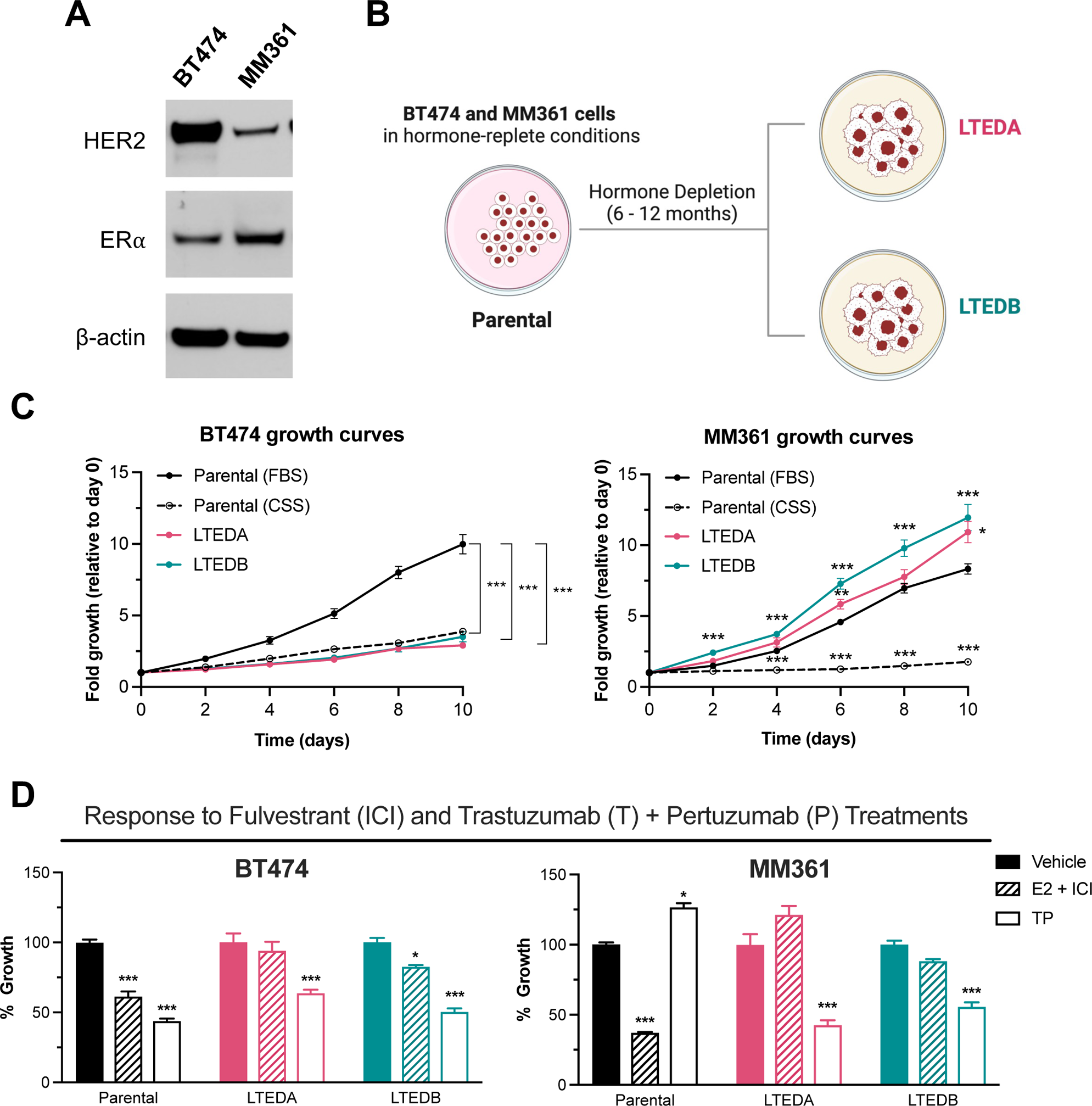
Growth pattern and response to ER- and HER2-targeted therapies of HER2+/ER+ LTED variants differ from their parental counterparts. **(A)** Western blot validation that BT474 and MM361 express HER2 and ER. **(B)** Schematic illustrates our preclinical modeling of ER-targeted therapy resistance by establishing two LTED variants (A and B) for each of the BT474 and MM361 cell lines. **(C)** Growth curves for parental (BT474 and MM361) and their LTED-derived (A and B) variants. Parental cells were cultured in either normal growth media (IMEM supplemented with 10% FBS) or hormone-deprived media (phenol-red free IMEM supplemented with 10% CSS). Parental cells of the CSS group were hormone-deprived for 72 h before seeding. Plots represent mean ± SEM from two independent experiments, each performed in triplicates and asterisks denote significant changes using one-way ANOVA followed by Dunnett’s multiple comparison test. **(D)** Parental MM361 is intrinsically resistant to TP treatments whereas both MM361 and BT474 LTED models are less responsive to ICI. Crystal violet growth assay after six days of treatments with either vehicle, 10 nM E2 + 1 µM ICI or 1 µg/ml T + 1 µg/ml P. Representative graph with data presented as meanL±LSEM from three independent experiments with six readings and analyzed by ANOVA using Dunnett’s multiple comparison test. \**p*L<L0.05, \*\**p*L<L0.01 and \*\*\**p*L<L0.001 were considered statistically significant.

Next, we evaluated our HER2+/ER+ parental/LTED pair growth response to HER2- and ER-targeted therapies: dual HER2 blockade by TP, and fulvestrant (ICI) as a selective estrogen receptor degrader (SERD). For BT474, LTED growth was significantly inhibited by TP and minimally inhibited by ICI, whereas both TP and ICI caused significant growth inhibition of parental cells (**Fig. 2D**, left panel). Compared to parental MM361, both MM361 LTED variants were also less responsive to ICI but more responsive to TP. Additionally, results from western blot analysis showed no difference in ER protein levels of BT474 and MM361 LTEDs upon treatment with ICI versus vehicle, supporting the loss of ICI-mediated growth inhibition in HER2+/HR+ ETR LTEDs (**Fig. S1A** and **S1B**, (28)). Altogether, data shows that our two HER2+/ER+ LTED models differ from their parental counterparts in base line growth and responses to HER2- and ER-targeted therapies.

### MM361, but not BT474, LTEDs lose ER expression and E2-induced ER target gene expression

We further characterized our HER2+/ER+ ETR models to determine if alterations in the basal expression of ER and HER family members may explain observed differential growth inhibition of parental/LTED pair models to anti-HER2 and ER treatments. When compared to their corresponding parentals, BT474 LTEDs showed no substantial changes in ER protein or *ESR1* transcript levels, whereas a significant reduction of both was observed in MM361 LTEDs (**Fig. 3A** to **3C**). Additionally, ER transcriptional activity was confirmed by increased expression of ER-target genes (*PGR* and *TFF1*) upon E2 stimulation only in LTEDs of BT474, but not MM361 (**Fig. 3D**). Protein analysis of the HER family revealed a significant upregulation of EGFR and a modest increase of HER3 in BT474 LTEDs (**Fig. 3A** and **3B**). A significant increase in HER2 and/or modest increase in EGFR (which is able to form heterodimers with HER2 (40)) protein levels of MM361 LTEDs (**Fig. 3A** and **3B**) may explain why these cells responded better to growth inhibition by TP than parentals (**Fig. 2D**). LTED variants of both BT474 and MM361 cell lines showed activation of the pro-survival AKT signaling (pAKT) downstream of HER2 (**Fig. 3A** and **3B**).

**Figure 3.**
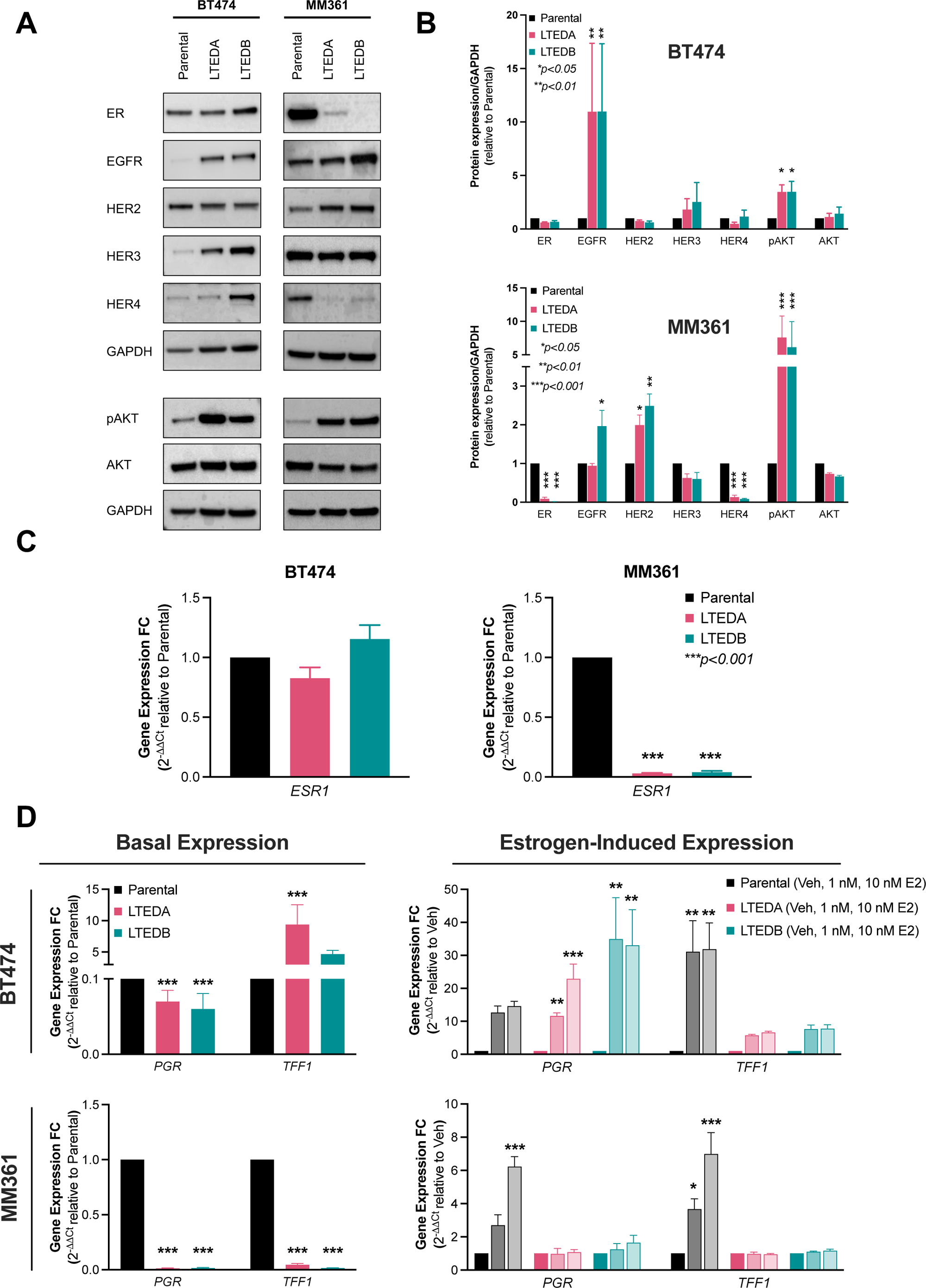
Expression and genomic activity of ER is either preserved or lost in our HER2+/ER+ LTED models. **(A)** Western blot analysis show that ER protein is drastically reduced in the MM361 LTED models but preserved in the BT474 LTEDs. **(B)** Blots from (A) were analyzed by densitometry using ImageJ. Protein expressions were normalized to GAPDH and graphed as meanL±LSEM from at least two independent experiments. ANOVA followed by Dunnett’s multiple comparison test was performed to compare groups and statistical significance is shown on the graph. **(C-D)** Expression of *ESR1* and ER-target genes (*PGR* and *TFF1*) are maintained in BT474 LTEDs but drastically reduced in MM361 LTEDs. mRNA levels of ER and selected ER-target genes under basal conditions or after estrogen stimulation (with either 1 nM or 10 nM E2 for 72 h) were analyzed by qRT-PCR and data represented as mean ± SEM of fold changes in transcript expression from four independent experiments. Asterisks denote significant changes using ANOVA followed by Dunnett’s multiple comparison test. \**p* < 0.05, \*\**p* < 0.01 and \*\*\**p* < 0.001 were considered statistically significant.

### MM361 LTEDs gain basal-like and HER2-enriched intrinsic subtypes and exonic mutations in genes encoding transcription factors and chromatin modifiers

Resistance to ET may arise due to changes in gene transcription and/or mutational alterations. The loss/downregulation of ER and increase in HER2 expression in the MM361 LTED model (**Fig. 3**), coupled with their more rapid adaptation to LTED conditions and increased growth rate vs. parentals (**Fig. 2C**), raised the question of whether this model has shifted its intrinsic subtypes. Data from the SONABRE registry study supports this notion, reporting that HER2+/ER+ BCa has the highest rate of subtype discordance at metastatic presentation (41). We therefore prioritized the MM361 LTED model for further transcriptomic and genomic analyses to identify putative molecular mechanisms driving resistance. We first performed scRNAseq and analyzed the molecular subtype of individual cells using PAM50. MM361 LTEDs intrinsic subtyping is discordant from its parentals. As might be predicted from their loss/downregulation of ER expression at the mRNA and protein levels, LTEDs demonstrated a shift towards non-luminal HER2-enriched (HER2-E) and basal-like phenotypes (**Fig. 4A** and **4B**). Additionally, we performed whole-genome sequencing to identify mutations gained in MM361 LTED. Analysis of SBS mutational signatures from our WGS data showed an enrichment in C to T base substitutions (SBS1) as well as T to C base substitutions (SBS5) across the genome of MM361 LTEDs compared to the parental cell line (**Fig. 4C**). These aging-related, clock-like mutational signatures are prevalent in many cancer types, though SBS1 mutations were recently reported to be enriched in breast cancer metastases in an age-independent manner (42). Next, we focused on mutated genes at their exonic regions. MM361 LTEDA and B shared a total of 70 genes bearing mutations in their exonic regions when compared to their parental cells (**Fig. 4D**). After excluding the 22 genes with silent mutations and characterizing the mutational signatures of the remaining 48, C to T and C to A were the most predominant mutation types (**Fig. 4E** and **Table S2** (28)). Most of these mutated genes were modestly deleterious (**Fig. 4F**) as predicted by Varmap (43). These genes mainly encode for transcription factors and chromatin modifiers, namely *HEY1*, *CHD4*, *MAFF*, *PRDM14*, *SATB2*, *SUPT6H*, *ZNF135*. Importantly, mutation of one or more of these genes is significantly enriched in ETR advanced breast cancers that are HER2+ and/or progesterone receptor-negative (PR-; **Fig. S2A** and **S2B** (28)). Together, these data show that a shift towards more aggressive intrinsic molecular subtypes (basal and HER2-enriched) and acquiring mutated transcription factors and chromatin modifiers accompanies the development of ETR in the MM361 LTED model.

**Figure 4.**
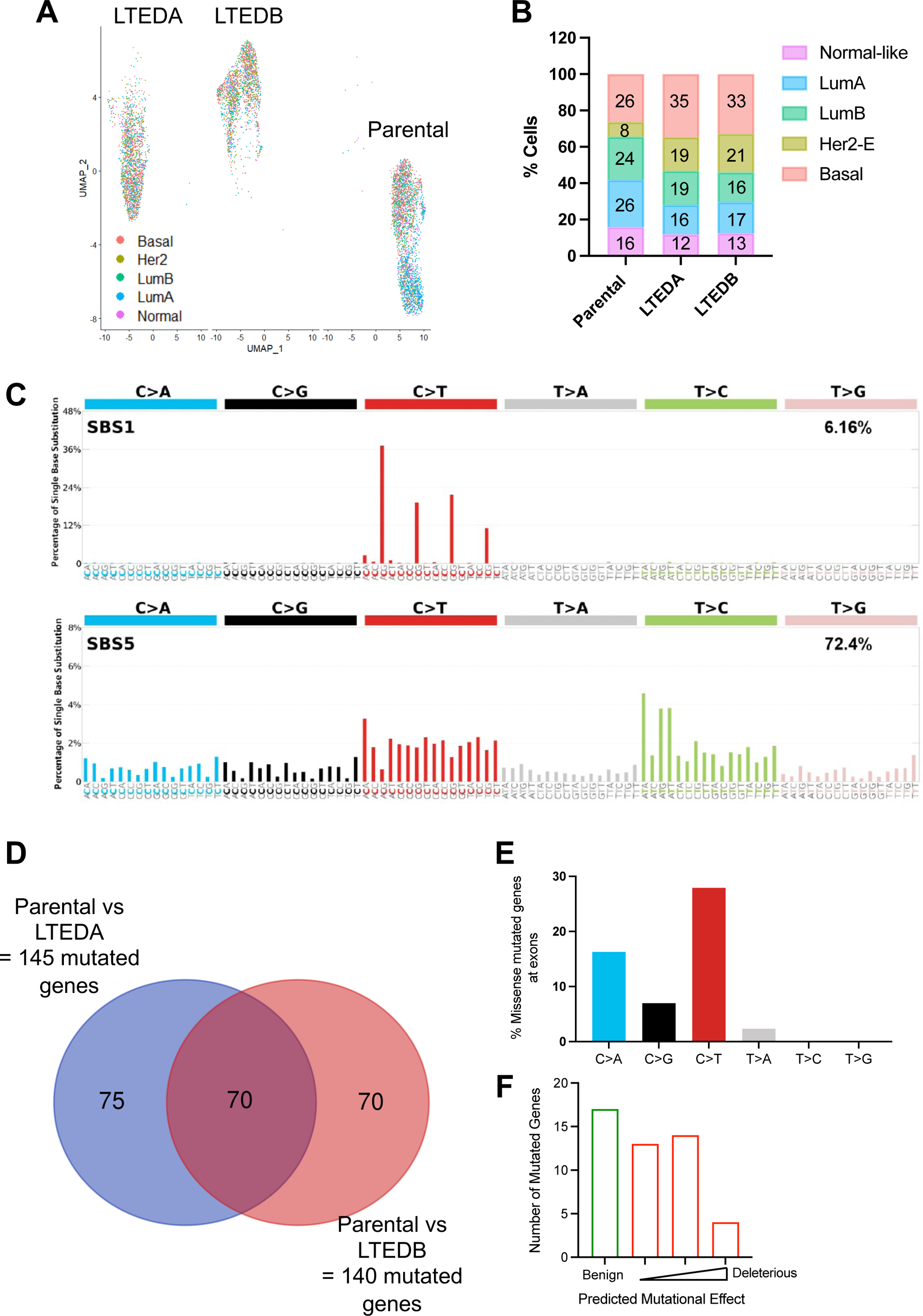
MM361 LTEDs gain basal-like, HER2-E intrinsic subtypes and harbor mutations in transcriptional and chromatin regulatory factors. **(A)** UMAP plot of scRNAseq data grouped by predefined PAM50 intrinsic molecular subtypes. **(B)** Bar graph quantifies the distribution of intrinsic subtypes. **(C-F)** Characterizing MM361 LTED mutations using WGS. **(C)** Single base substitution (SBS) mutational signatures for the entire genome of MM361 LTEDs. **(D)** Venn diagram intersection showing shared mutated genes at their exonic regions of MM361 LTEDA and B versus parental cells. **(E)** Quantifying SBS for non-silent mutated genes at exonic regions. **(F)** Mutational effect predictions using Varmap (https://www.ebi.ac.uk/thornton-srv/databases/VarMap, accessed on 11 August 2022).

### Lipid metabolism pathways are upregulated in the MM361 LTEDs

Using our scRNAseq data, we performed unsupervised clustering using Seurat to identify over- and under-represented cell populations in MM361 parental and LTED A and B cells (**Fig. S3** (28)). We identified seven clusters, with clusters 2 and 5 enriched in parental cells and clusters 0 and 6 enriched in LTED cells (**Fig. S3A** to **S3C** (28)). GSEA analysis showed that parental-dominant clusters 2 and 5 were enriched with genes regulating estrogen response, glycolysis, TCA cycle, and oxidative phosphorylation (**Fig. S3D** (28)). By contrast, LTED-dominant clusters showed a significant overrepresentation of fatty acid metabolism (in the more abundant cluster 0), and RNA processing pathways e.g., nonsense-mediated decay (in the less abundant cluster 6). We then constructed “pseudo-bulk” profiles from scRNAseq and performed differential expression analysis of transcriptomic changes that characterize MM361 LTEDA and B vs. parental cells. Transcriptional profiling at a false discovery rate (FDR) <0.05 and a log2 fold change ≤-0.5 and ≥0.5 identified a total of 356 differentially expressed genes (DEG) between parental and LTEDA, and 674 DEG between parental and LTEDB (**Fig. 5A**). The intersection of DEG from LTED vs. parental comparisons resulted in 202 DEG between both LTED lines and parental cells, with 156 and 46 genes being upregulated and downregulated, respectively (**Fig. 5B** and **Table S3** (28)). We validated the transcriptomic analysis of MM361 LTEDs and confirmed the increased expression of eleven upregulated DEGs using qRT-PCR (**Fig. 5C**). Importantly, *ESR1* and *TFF1* were among the top downregulated genes and *ERBB2* expression was upregulated (**Fig. S4** (28)), consistent with data shown in **Fig. 3**. Using LISA (44) to predict the transcriptional regulators (TRs) of these DEGs (**Table S3** (28)), we found that the steroid hormone receptors *PGR*, *ESR1*, and *AR*, as well as the pioneer factors *GRHL2* and *FOXA1* were among the most enriched TRs in common for both up- and downregulated DEGs. Predicted TRs associated exclusively with downregulated DEGs included *E2F1*, *RARA*, and *MAFB* (a homolog of *MAFF*, which we identified as mutated in LTED cells, **Table S2** (28)). Predicted TRs for upregulated DEGs included *GATA2*, *PPARG*, and *CEBPA*.

**Figure 5.**
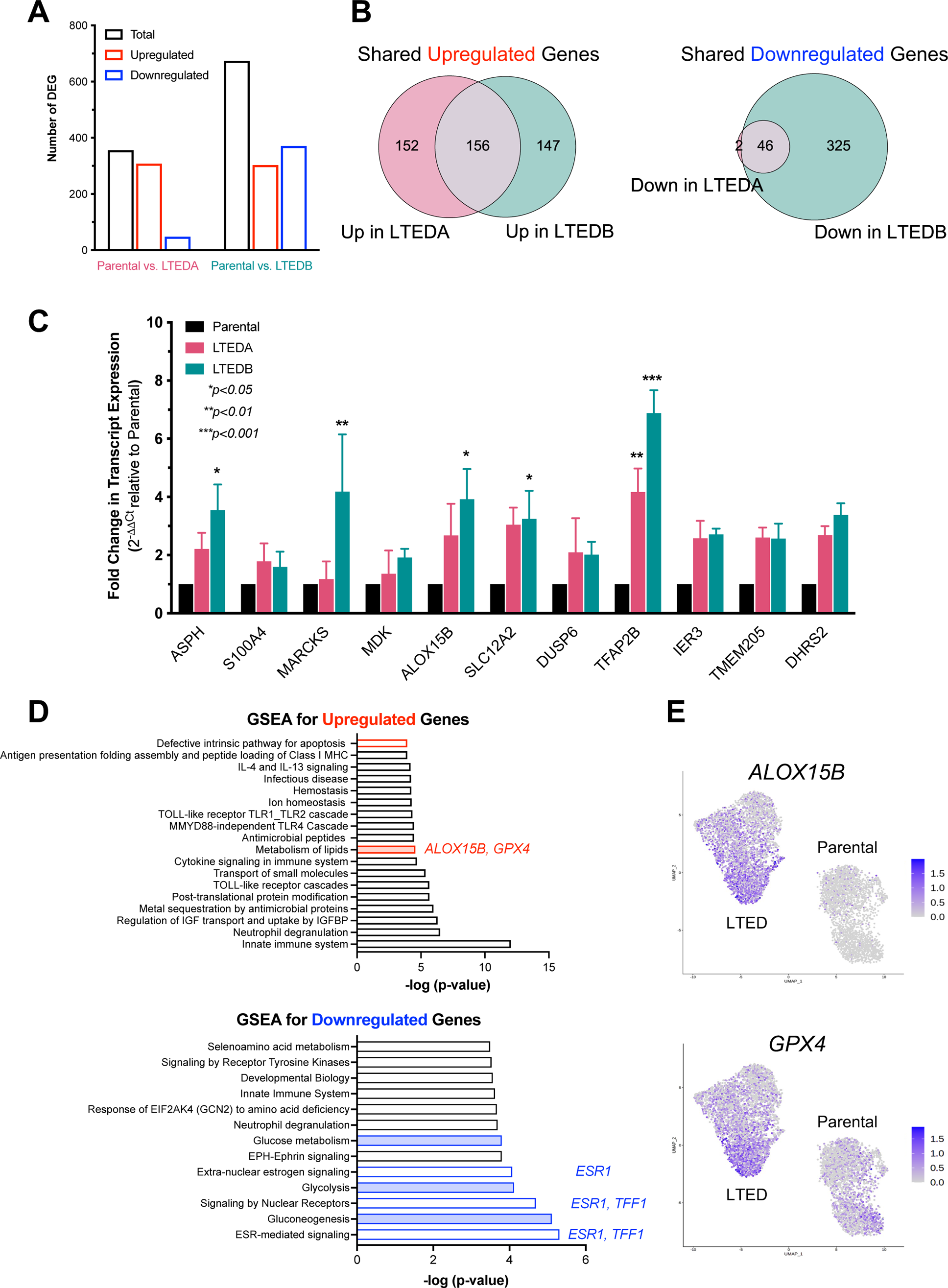
Lipid metabolism pathways are upregulated in LTEDs of the MM361 cell line. **(A)** Transcriptional profiling from scRNAseq at FDR<0.05 and log2 fold change ≤-0.5 and ≥0.5 identified 356 DEG between parental and LTEDA and 674 DEG between parental and LTEDB. **(B)** A total of 202 DEG are shared in both LTED lines vs. parental cells. Venn diagram intersections show 156 shared upregulated and 46 shared downregulated DEG in the MM361 LTEDs vs. parental. **(C)** qRT-PCR was used to confirm increased expression of selected upregulated genes identified from the scRNAseq transcriptomic profiling. Data presented as mean ± SEM of fold changes in expression of each gene (LTEDs relative to parental cells) from three independent experiments. Asterisks denote significant changes using ANOVA followed by Dunnett’s multiple comparison test. **(D)** Top 20 enriched biological processes in MM361 LTEDs. Overlapped up- and down-regulated DEGs (from B) in MM361 LTEDs were analyzed by GSEA using the REACTOME gene set, with FDR q-values <0.05. Red- and blue-filled boxes highlight metabolic pathways. **(E)** UMAP plots for *ALOX15B* and *GPX4,* among the upregulated genes of the lipid metabolism pathway being enriched in the MM361 LTEDs.

To identify biological processes that are enriched within the LTED variants, we performed GSEA on the upregulated and downregulated shared gene lists of both LTED lines vs. parental cells. Genes upregulated in LTEDs were enriched in pathways involved in defective apoptosis and immune-related pathways. Conversely, pathways involved in estrogen and other nuclear receptor signaling were enriched within the downregulated gene set (**Fig. 5D**). Notably, metabolic pathways were among the enriched pathways in the MM361 LTEDs. While lipid metabolism was upregulated, pathways of glucose metabolism, glycolysis and gluconeogenesis were all downregulated (**Fig. 5D**), consistent with our unsupervised clustering analysis results (**Fig. S3** (28)). The upregulation of lipid metabolism pathways is also supported by the observed enrichment of putative TRs *PPARG* and *CEBPA* from our upregulated DEGs (**Table S3** (28)), which are key activators of lipid metabolism (45). Our data suggest notable metabolic remodeling of MM361 LTEDs, specifically upregulation of lipid metabolism.

### Dual targeting of HER2 and GPX4 increase cell death of HER2+/ER+ LTEDs

*ALOX15B* and *GPX4* are two of the upregulated genes of lipid metabolism pathways enriched in the MM361 LTEDs (**Fig. 5D** and **5E**). In addition, *ALOX15B* and *GPX4* were among the 13 MM361 LTED DEGs that overlapped with ferroptosis-related genes (**Fig. S5** (28)). Ferroptosis is an iron-dependent cell death that is triggered by lipid peroxidation (46). Interestingly, both upregulated proteins are opposing key players in regulating ferroptosis (47); ALOX15B induces ferroptosis, whereas GPX4 inhibits it. We validated the upregulation of *ALOX15B* mRNA in MM361 LTEDs (**Fig. 5C**), and GPX4 protein levels in both MM361 and BT474 LTEDs (**Fig. 6A** and **S6A** (28)).

**Figure 6.**
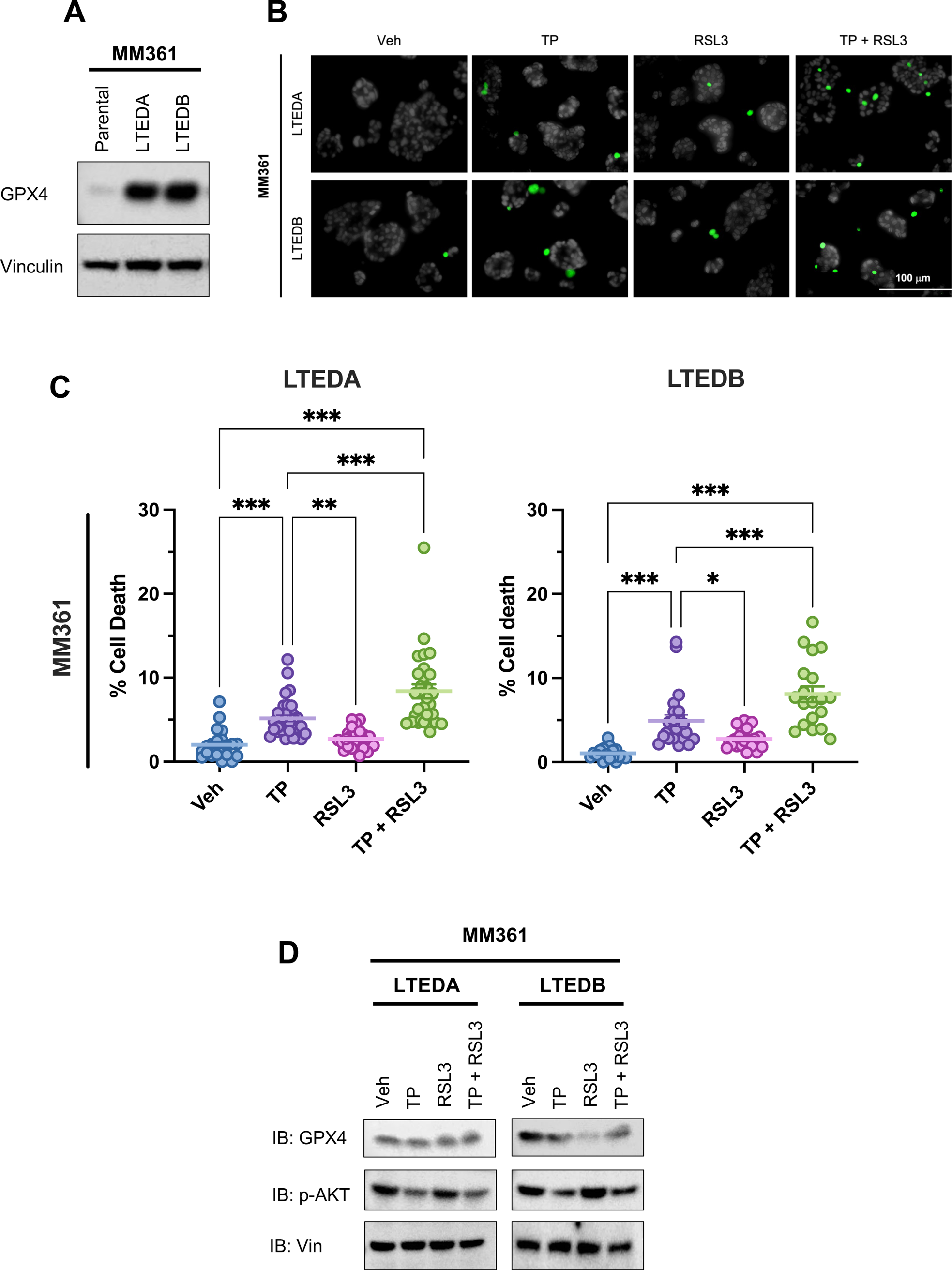
Dual targeting of HER2 and GPX4 increase cell death of MM361 LTEDs. **(A)** Representative immunoblot of three independent experiments showing enhanced expression of GPX4 protein in MM361 LTEDs. **(B-D)** Cells treated with either vehicle, 1 µg/ml T + 1 µg/ml P, 1LμM RSL3 or a combination of both for 72 h then stained with Hoechst to monitor total cell number (blue nuclei), and Sytox green to monitor dead cells (green nuclei) as in panels B and C or analyzed by western blot analysis as in D. **(B-C)** Sytox staining experiments show that cotreatments of TP + RSL3 induce the highest cell death in MM361 LTEDs. **(B)** Cell death representative images and **(C)** quantifications from two independent experiments with at least ten fields analyzed for each and data presented as meanL±LSEM of cell death percentages. Asterisks denote significant changes using one-way ANOVA followed by Tukey’s multiple comparison test; \**p*L<L0.05 and \*\*\**p*L<L0.001 were considered statistically significant. **(D)** Treatment effects on GPX4 and pAKT protein levels were assessed with western blot analysis. Immunoblots are a representation of two independent experiments.

Inhibiting the PI3K-AKT-mTOR signaling pathway – which we show in **Fig. 3** is strongly activated in both MM361 and BT474 LTED cells – can sensitize BCa cells to ferroptosis induction, in part by reducing SREBP-mediated lipogenesis (48). Therefore, we hypothesized that LTED cells of HER2+/ER+ BCa are resistant to ferroptosis, due to GPX4 upregulation, and that dual targeting of HER2 (upstream of PI3K-AKT-mTOR) and GPX4 may sensitize these LTEDs to ferroptosis-induced cell death. In MM361 LTEDs, significant cell death was observed following HER2 inhibition by TP, but not GPX4 inhibition by RSL3 (**Fig. 6B** and **6C**). In BT474 LTEDs, TP and RSL3 alone each significantly increased cell death (**Fig. S6B** and **S6C** (28)). Importantly, in both LTED models, cell death was significantly enhanced by the combination of TP + RSL3. As expected, TP alone reduced the activation of AKT (pAKT) and RSL3 alone reduced protein levels of GPX4 (**Fig. 6D**), but no further reduction was observed with TP + RSL3 vs. that of the single treatments (**Fig. 6D**). To test whether the increased cell death induced by the TP + RSL3 combination was due to ferroptosis, we measured the levels of two ferroptosis lipid peroxidation markers, 4-hydroxynonenal (4-HNE) and malondialdehyde (MDA) (**Fig. S7** (28)). RSL3 alone slightly increased levels of both 4-HNE and MDA, but TP + RSL3 did not further enhance the levels of 4-HNE and MDA protein conjugates. Together, these data suggest that increased lipid metabolism in HER2+/ER+ BCa LTED models is accompanied by reprogramming of ferroptosis pathways, and that the combination of anti-HER2 agents and a ferroptosis inducer is an effective approach to inducing cell death.

## DISCUSSION

To date, models of acquired ETR have been almost exclusively developed for HER2−/ER+ BCa, and not HER2+/ER+. In this study, we generated and characterized models of the latter without using genetic manipulation approaches. We report that ETR BT474 and MM361 AI-resistant (LTED) models capture distinct phenotypes of HER2+/ER+ BCa as supported by the following results. Compared to parentals, MM361 LTEDs grew faster, lost ER, and increased HER2 expression, whereas BT474 LTEDs grew slower and showed no substantial changes in ER or HER2 expression. Additionally, growth inhibition by anti-HER2 (TP) treatment was much more pronounced in the LTEDs vs. parental MM361, unlike the BT474 model, where both LTEDs and parentals benefited from TP. Differences in ER and HER2 expression in the LTEDs of MM361 and BT474 models may suggest distinct resistance mechanisms to ET partly due to the differential distribution of intrinsic subtypes within these two cell lines. It could imply that BT474 ETR behaves more like luminal whereas MM361 LTEDs behaves more like non-luminal HER2+ BCa and hence the more aggressive nature of the latter.

HR and HER2 receptor expressions, or lack thereof, are critical in guiding BCa treatments. However, pathology-based immunohistochemistry (IHC) does not fully recapitulate intrinsic biology. Instead, substantial discrepancies exist between IHC-based and gene expression-based PAM50 subtyping (49). Our results show that MM361 LTEDs intrinsic subtyping is discordant from its parentals. We observed a loss in ER/*ESR1* accompanied by a gain in HER2/*ERBB2* expression and, thus, molecular switching to non-luminal HER2-E phenotype and sustained activation of AKT signaling. These alterations altogether explain why our ETR MM361 LTEDs were less responsive to ICI but more responsive to anti-HER2 targeted therapies vs. their corresponding TP-resistant parentals. HER2-E subtype represents ∼75% HER2+/HR- and 30% of HER2+/HR+ tumors (49,50), is associated with anti-HER2 sensitivity (49), and is a predictive biomarker of poor response and resistance to AI (10). Our findings are supported by several studies. First, data from the SONABRE registry (NCT03577197) show that among all BCa subtypes, patients with HR+/HER2+ advanced BC exhibit the highest receptor subtype discordance rate between primary tumor and metastatic lesions as they convert mainly to HR+/HER2- and HR−/HER2+ (41). In other words, ER and HER2 expressions varied during tumor progression, and cross-talk between ER and HER2 was proposed as the leading cause of this discordance (41). Second, HER2 and ER expressions are negatively correlated in HER2+ BCa tumors (51). Third, cell lineage tracing experiments in mice showed that HER2+/ER+ cells can lose ER expression and be converted into highly aggressive HER2+/ER-BCa cells (52). Lastly, HER2 overexpression (with or without EGFR) activates downstream AKT and MAPK pathways to circumvent ER inhibition and promotes growth of ETR BCa (2,25,53). ER loss in our MM361 ETR model could be explained by *ESR1* promoter methylation and/or decreased FOXA1 and GATA3, which are required for normal ER expression (52). Additionally, losing the ER-regulated cluster of let-7c, miR99a, and miR125b (54), could explain our upregulated HER2 expression.

Beyond the HER2-E intrinsic subtype, acquiring mutations in genes encoding for transcription factors and chromatin modifiers could underlie mechanisms of AI resistance and/or the aggressive nature of the MM361 LTED model. For example, integrated genomic analysis of responding vs. resistant ER+ breast tumors from a patient who developed resistance to letrozole treatment revealed an acquired mutation in the *CHD4* gene (55). SUPT6H protein levels decrease in poorly differentiated breast tumors, and SUPT6H is a requirement for ERα transcriptional activity and maintenance of chromatin structure in BCa cells (56). Also, SATB2 induces transformation of mammary epithelial cells into progenitor-like cells and its knockdown attenuates proliferation and epithelial-mesenchymal transition (EMT) of BCa cells (57). What role the mutations in these genes that we observe in MM361 LTEDs play in the ETR phenotype is a subject of future study.

BCa cells carrying *PIK3CA* activating mutation (like the parental BT474 and MM361 cells (58)), and thus an activated PI3K-AKT-mTOR signaling pathway, are resistant to ferroptosis induction by the GPX4 inhibitor RSL3 (48). Consistent with this, inhibiting an activated PI3K-AKT-mTOR signaling pathway sensitizes BCa cells to ferroptosis induction by RSL3 (48). In the current study, we show that dual targeting of HER2 and GPX4 increases cell death of HER2+/ER+ LTEDs but not levels of ferroptosis lipid peroxidation markers beyond that of single-agent treatments. One possible explanation could be that GPX4 inhibition may not be sufficient to overcome ferroptosis resistance in our LTEDs. In other words, elevated GPX4 may not be the sole cause of ferroptosis resistance in MM361 LTEDs; instead, several other ferroptosis-related DEGs may mediate this resistance. Dysregulated oxidative and iron homeostasis exemplified by upregulation of ferroptosis suppressor genes (*TMBIM4* and *GPX4*), antioxidants (*PRDX2*), iron chelators (*LCN2*), and/or downregulation of ferroptosis inducing genes (*RPL8* and *HILPDA*) could be responsible for ferroptosis resistance in our LTEDs (**Fig. S3** (28)). Moreover, multiple ferroptosis regulatory pathways that are independent of GPX4 have recently been discovered (59). For example, cisplatin-resistant-derived exosomes secrete microRNAs that increase the expression of ferroptosis suppressor protein 1 (FSP1) and thus enhance resistance of cancer cells to ferroptosis (60). Along the same lines, LTED cells treated with TP + RSL3 may protect themselves from ferroptosis using a similar mechanism. In our hands, TP did not reduce GPX4 nor enhance expression of ferroptosis lipid peroxidation markers. However, a recent study reported that trastuzumab triggers ferroptosis by reducing GPX4 whilst increasing ROS levels in embryonic rat myoblast (H9c2) cells (61). This inconsistency could be attributed to variances in cell context, doses and/or duration of treatments.

Inevitably, our study has several limitations that must be considered. Future xenograft studies will be necessary to verify and follow up on some of our current findings, for example, the response of ETR HER2+/ER+ to ET and anti-HER2 treatments and whether ferroptosis induction combined with anti-HER2 treatments suppresses the growth of these cells *in vivo*. Despite harboring a sufficient gain in *ERBB2* copy number to be classified as HER2+ (amplified, (62,63)), a previous study demonstrated that the parental MM361 cell line exhibits heterogenous expression of *ERBB2* and *ESR1*, and this heterogeneity is not due to heritable genetic differences (63), suggesting a high degree of plasticity and variability. This informed our rationale for performing scRNAseq to capture transcriptional heterogeneity of the parental cells and ETR LTED models. Importantly, our conventional and “pseudo-bulk” analyses of the scRNAseq data both identified alterations in lipid metabolism. However, we sequenced a relatively small number of cells by scRNAseq (<4000 cells per sample), and combined with the fact that scRNAseq is best suited to detecting higher abundance transcripts, we cannot exclude the possibility that we are missing minor populations or subclones that contribute significantly to the ETR phenotype (e.g. (64,65)). The gene mutations we identified in LTEDs are constrained by the WGS coverage that we achieved (LTEDA: 53x, LTEDB: 50x, Parental: 52x). Furthermore, deeper sequencing depths may capture and result in a larger number of mutated genes. However, even at the sequencing depths and coverage levels we used, our results may suggest that multiple potential resistance mechanisms exist. Deep sequencing of a different epithelial tumor, colorectal cancer, coupled with an evolutionary analysis, showed that the ∼10^9^ cells in a 1 cm^3^ lesion will contain every possible resistance mutation in a minor subclone within the lesion (66). This large diversity of resistance mechanisms implies a need for treatment regimens that include therapies too numerous to be given simultaneously in combination (67).

Increasing efforts are being directed toward chemotherapy-sparing regimens for HER2+/ER+ BCa (68–71). However, these treatment strategies should also be tested on HER2+/ER+ BCa that are refractory to or have progressed on ET. Therefore, future studies are needed to address open questions regarding 1) responsiveness of ETR HER2+/ER+ models to dual and triple combinations of anti-HER2, ET/SERDs (including recently approved oral SERD amcenestrant), and CDK4/6 inhibitors, as well as innovative sequencing of these combinations (63,64,67,72), 2) the impact of our identified mutations on development of ETR in HER2+/ER+ BCa, and 3) differences in ER activity between ETR models of HER2+/ER+ (as BT474) and other HER2-ER+.

## CONCLUSIONS

Characterizing models of HER2+/ER+ that mimic a real-world treatment pattern is necessary to understand mechanisms of resistance to ET and may provide useful information for refining current treatment approaches and improving patients’ outcomes. Here, we report that anti-HER2 targeted therapies effectively inhibit growth of ETR HER2+/ER+ BCa cells that exhibit concurrent loss of ER expression and gain in HER2 and HER2-E phenotype. Our BT474 and MM361 AI-resistant models capture distinct phenotypes of HER2+/ER+ BCa and pinpoint altered lipid metabolism and ferroptosis remodeling as vulnerabilities of this type of ETR BCa.

## ABBREVIATIONS

4-HNE: 4-hydroxynonenal
BCa: Breast cancer
BT474: BT-474
CSS: Charcoal-stripped bovine serum
DEG: Differential expressed gene
DMFS: Distant metastasis-free survival
E2: 17β-estradiol
EMT: epithelial-mesenchymal transition
ER: Estrogen receptor
ET: Endocrine therapy
ETR: Endocrine therapy-resistant
FDR: False discovery rate
FSP1: Ferroptosis suppressor protein 1
GSEA: Gene set enrichment assay
HER2: human epidermal growth factor receptor 2
HR: Hormone receptor
ICI: fulvestrant
IMEM: Improved minimum essential medium
LTED: Long-term estrogen deprivation
MDA: Malondialdehyde
MM361: MDA-MB-361
MSigDB: Molecular signature database
OS: Overall survival
PFS: Progression-free survival
qRT-PCR: Real-time PCR
RFS: Relapse-free survival
SBS: Single base substitution
scRNAseq: Single-cell RNA sequencing
SERD: Selective estrogen receptor degrader
TR: Transcriptional regulator
TP: Trastuzumab + pertuzumab
WGS: whole genome sequencing

## Acknowledgements

The authors would like to thank members of the Riggins laboratory, Drs. Michael Johnson, Marc Lippman, and Joyce Slingerland (Lombardi Comprehensive Cancer Center, Georgetown University), and members of NR IMPACT for sharing reagents, scientific insights, and/or technical assistance.

## Data Availability

Original data generated and analyzed during this study are included in this published article or in the data repository listed in References ((28); https://doi.org/10.5281/zenodo.8265999).

## Authors’ contributions

SB, HS and RBR designed the experiments. SB performed most of the experiments, analyzed data, prepared figures, and drafted the manuscript. HS performed, analyzed, and graphed some of the RT-PCR and growth assays. MB and MP performed the scRNAseq experiments and LJ analyzed the scRNAseq and WGS data. ST and DM handled most of the western blot analyses. MDM and RAB contributed to the study discussions. RBR supervised the study, wrote parts of the manuscript, reviewed, and revised it. All authors read, revised, and approved the final manuscript.

## Ethics approval and consent to participate

Not applicable.

## Consent for publication

All authors agreed to publish this study.

## ADDITIONAL FILES

**Additional file 1:** Supplemental materials and methods. **Table S1.** List of primer sequences used for the detection of transcripts. Supplemental figures and figure legends: **Figure S1.** Effects of HER2- and ER-targeted therapies on ER expression and phosphorylation of HER2 in HER2+/ER+ LTEDs. **Figure S2.** Breast tumors harboring mutated genes identified in MM361 LTED cells are associated with HER2+ status and lower PR expression. **Figure S3.** Cluster analysis of MM361 parental and LTED cell scRNAseq data. **Figure S4.** scRNAseq expression of *ESR1* and *ERBB2* in MM361 parental and LTED cells. **Figure S5.** Ferroptosis-related genes in MM361 LTEDs. **Figure S6.** Dual targeting of HER2 and GPX4 increase cell death of BT474 LTEDs. **Figure S7.** Effects of TP and RSL3 on protein expression of ferroptosis lipid peroxidation markers (4-HNE and MDA).

**Additional file 2: Table S2.** Shared genes bearing exonic mutations in MM361 LTEDA and B versus parental cells and predicted pathogenicity.

**Additional file 3: Table S3.** Differentially expressed genes that are shared in the LTEDA and LTEDB variants of the MM361 cell line, and inference of transcriptional regulators using LISA (http://lisa.cistrome.org/, accessed on 4 May 2023).

