## Additional File 1, Supplemental Figures for "Unraveling Vulnerabilities in Endocrine Therapy-Resistant HER2+/ER+ Breast Cancer"

#### SUPPLEMENTAL MATERIALS and METHODS

**Table S1:** List of primer sequences used for the detection of transcripts.

| Gene | Primer Sequence | Reference |
|---|---|---|
| **ESR1** | **Forward:** 5’-AGAGGGCATGGTGGAGATCTT-3’<br>**Reverse:** 5’-CAAACTCCTCTCCCTGCAGATT-3’ | (1) |
| **PGR** | **Forward:** 5’-CAGTGGGCAGATGCTGTATT-3’<br>**Reverse:** 5’-GGATCTGCCACATGGTAAGG-3’ | |
| **TIFF1** | **Forward**: 5’-CCCTCCCAGTGTGCAAATAA-3’<br>**Reverse:** 5’-GATCCCTGCAGAAGTGTCTAAA-3’ | |
| **ASPH** | **Forward:** 5’-CATGGAGGACACAAGAATGGG-3’<br>**Reverse:** 5’-CCAAACGACAGCTACAGATGT-3’ | (2) |
| **S100A4** | **Forward:** 5’-AGTTCAAGCTCAACAAGTCAGAACTAA-3’<br>**Reverse:** 5’-TCATCTGTCCTTTTCCCCAAGA-3’ | (3) |
| **MARCKS** | **Forward:** 5’-TCTCCTGTCCGTTCGCTTTG-3'<br>**Reverse:** 5’-CCAGTTCTCCAAGACCGCAG-3' | (4) |
| **MDK** | **Forward:** 5’-ATGCAGCACCGAGGCTTCCT-3'<br>**Reverse:** 5’-ATCCAGGCTTGGCGTCTAGT-3' | (5) |
| **ALOX15B** | **Forward:** 5’-CAATGCCGAGTTCTCCTTCCATG-3'<br>**Reverse:** 5’-TGATGTGCAGGGTGTATCGGGT-3' | (6) |
| **SLC12A2** | **Forward:** 5’-ATCAATTTTTCAGTATTCCATGCATC-3'<br>**Reverse:** 5’-ACGCCATCCTGGAGATTTTG-3' | (7) |
| **DUSP6** | **Forward:** 5’-AACAGGGTTCCAGCACAGCAG-3'<br>**Reverse:** 5’-GGCCAGACACATTCCAGCAA-3’ | (8) |

| | | |
|---|---|---|
| **TFAP2B** | **Forward:** 5’-TGAAGATGCCAATAACAGCGGCA-3’<br>**Reverse:** 5’-GGAGCAAAACACCTCGCCGGT-3’ | (9) |
| **IER3** | **Forward:** 5’-AGTCTGGTGGTGGGTCGTAAGT-3’<br>**Reverse:** 5’-GATGGAAGGATCTCACGGATCT-3’ | (10) |
| **TMEM205** | **Forward:** 5’-GCAAATGTGGGTGACCTTCGTC-3’<br>**Reverse:** 5’-GCAGAGGTTGATGAAGGCACAG-3’ | |
| **DHRS2** | **Forward:** 5’-GGTGTCTACAATGTCAGCAAGA-3’<br>**Reverse:** 5’CACGCAGTTTACCCGGATGT-3’ | |
| **ACTB** | **Forward:** 5’-AAGCCACCCCACTTCTCTCTAA-3’<br>**Reverse:** 5’-CACCTCCCCTGTGTGGACTT-3’ | (11) |

**SUPPLEMENTAL FIGURES and FIGURE LEGENDS**

**Figure S1. Effects of HER2- and ER-targeted therapies on ER expression and phosphorylation of HER2 in HER2+/ER+ LTEDs. (A-B)** Western blot assessment of ER expression and phosphorylation of HER2 in BT474 LTEDs (A) and MM361 LTEDs (B) upon 6 h of treatment with either vehicle, 10 nM E2, 10 nM E2 + 100 nM ICI or 100 µg/ml T + 100 µg/ml P. Blots were analyzed by densitometry using ImageJ. Protein expressions were normalized to GAPDH and graphed as mean ± SEM from at least two independent experiments. One-way ANOVA followed by Dunnett's multiple comparison test was performed to compare groups and statistical significance is shown on the graph. The arrow on the MM361 LTEDB blot indicates correct molecular weight of ER (other bands are non-specific).

**Figure S2. Breast tumors harboring mutated genes identified in MM361 LTED cells are associated with HER2+ status and lower PR expression. (A-B)** Data were obtained from Razavi *et al*. (12) using cBioPortal for Cancer Genomics (http://www.cbioportal.org/, accessed on 11 May 2023)**.** Stacked bar charts illustrate HER2 status (classified by FISH and IHC; thus HER2+ indicates *ERBB2*/HER2 amplification) (A) and PR status (B) distributions in breast tumors with and without mutational alterations in at least one gene in which we identified non-silent exonic mutation in LTED cells (mutated gene list in Table S2). Proportions were tested by chi-squared analysis. Number of samples and statistical significance are shown on the graph.

**Figure S3. Cluster analysis of MM361 parental and LTED cell scRNAseq data. (A)** UMAP plot of scRNAseq data grouped by unsupervised clustering (10 Principal Components, Resolution 0.5) into seven clusters (0-6) across parental and LTED cell lines. **(B-C)** Sankey plot (B) and tabular (C) representation of the number of sequenced cells belonging to each cluster. **Bold**

clusters **0** and **6** are enriched in LTED cells, while *italicized* clusters *2* and *5* are enriched in parental MM361 cells. **(D)** Top enriched biological processes in parental- (2, 5) and LTED-dominant (0, 6) cell clusters, respectively. Genes from clusters were analyzed by GSEA using the Hallmark and REACTOME gene sets, with FDR q-values <0.05. Specific clusters in which these gene sets were enriched are indicated.

**Figure S4. scRNAseq expression of *ESR1* and *ERBB2* in MM361 parental and LTED cells.** UMAP plot of scRNAseq data shaded by expression of the indicated genes.

**Figure S5. Ferroptosis-related genes in MM361 LTEDs.** Analysis of the ferroptosis database (FerrDb V2) (13) and other literature search show that 13 of our DEGs are ferroptosis-related. Red and blue indicate upregulated and downregulated genes, respectively.

**Figure S6. Dual targeting of HER2 and GPX4 increase cell death of BT474 LTEDs. (A)** Representative immunoblot of three independent experiments showing enhanced expression of GPX4 protein in BT474 LTEDs. **(B-C)** Sytox staining experiments show that TP + RSL3 induces the highest cell death in BT474 LTEDs. Cells treated with either vehicle, 1 µg/ml TP, 250 nM RSL3 or a combination of both for 72 h then stained with Hoechst to monitor total cell number (gray nuclei), and Sytox green to monitor dead cells (green nuclei). (B) Cell death representative images and (C) quantifications from two independent experiments with at least 10 fields being analyzed for each. Data presented as mean ± SEM of cell death percentages using one-way ANOVA followed by Tukey's multiple comparison test. *$p < 0.05$ and ***$p < 0.001$ were considered statistically significant.

**Figure S7. Effects of TP and RSL3 on protein expression of ferroptosis lipid peroxidation markers (4-HNE and MDA).** MM361 LTED cells were treated with either vehicle, 1 µg/ml T + 1 µg/ml P, 1 µM RSL3 or a combination of both for 72 h then whole cell lysates were analyzed by western blot analysis. Immunoblots are a representation of two independent experiments. Blots were analyzed by densitometry using ImageJ and protein expression was normalized to vinculin.

Fig S1.

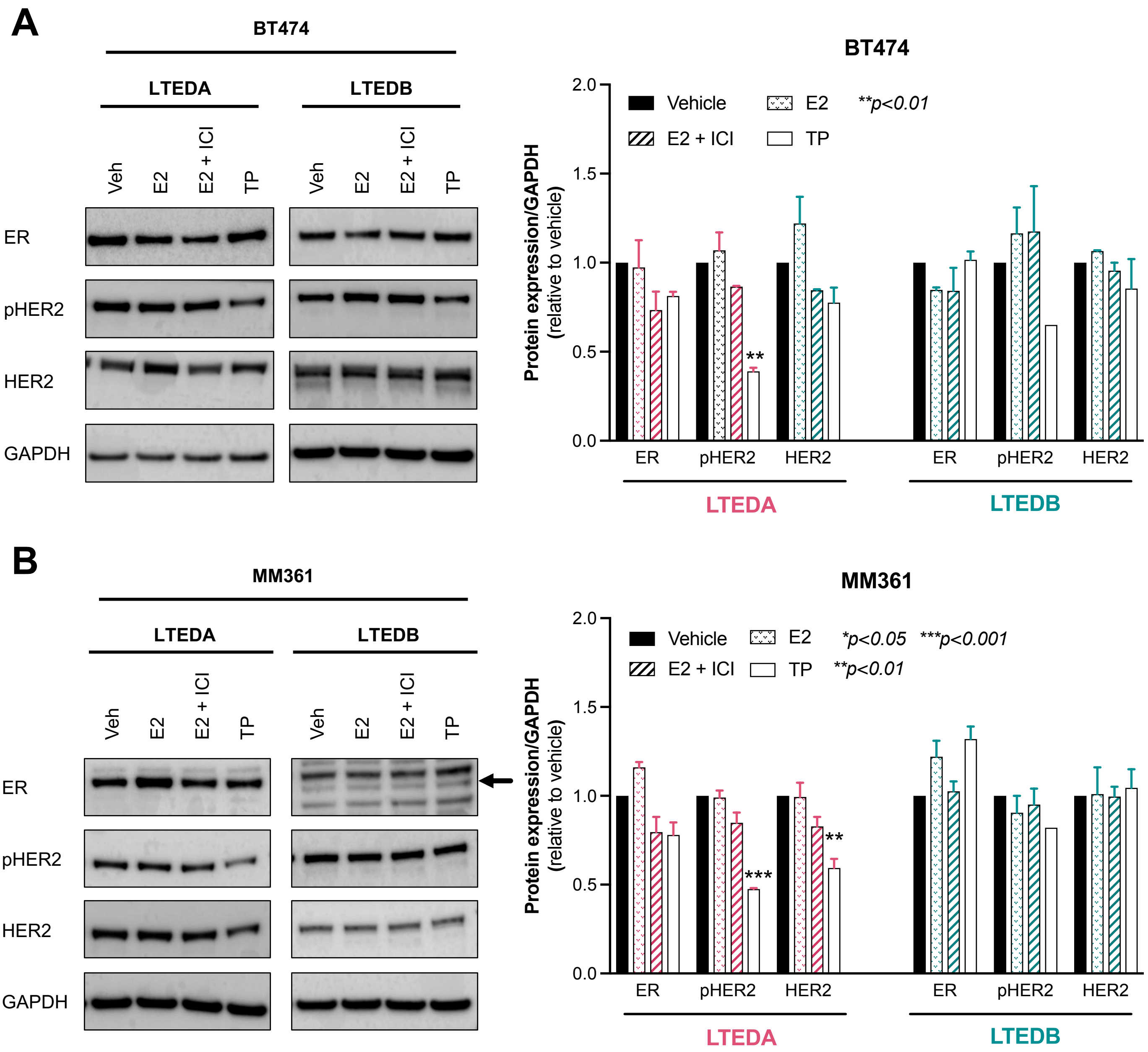
A
BT474
LTEDA
LTEDB
Veh
E2
E2 + ICI
TP
ER
pHER2
HER2
GAPDH
BT474
Protein expression/GAPDH (relative to vehicle)
Vehicle
E2
E2 + ICI
TP
**p<0.01
LTEDA
LTEDB
B
MM361
LTEDA
LTEDB
Veh
E2
E2 + ICI
TP
ER
pHER2
HER2
GAPDH
MM361
Protein expression/GAPDH (relative to vehicle)
Vehicle
E2
E2 + ICI
TP
*p<0.05 ***p<0.001
**p<0.01
LTEDA
LTEDB

**Figure S2.**

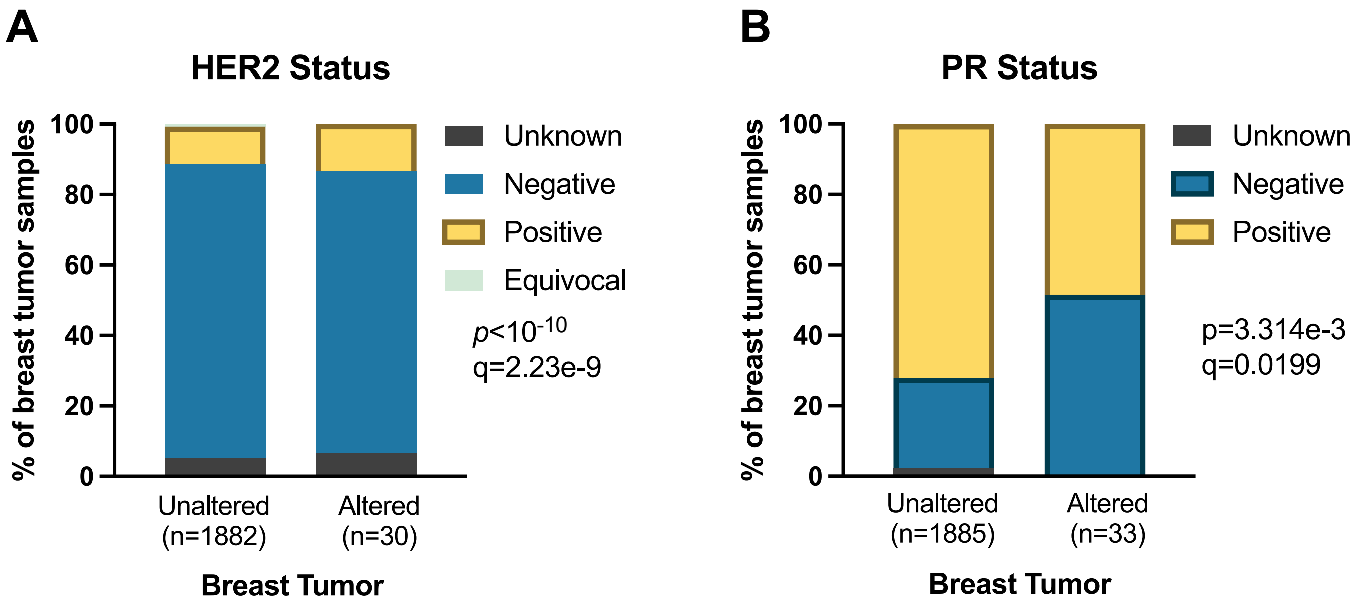
A
HER2 Status
% of breast tumor samples
100
80
60
40
20
0
Unknown
Negative
Positive
Equivocal
p<10-10
q=2.23e-9
Unaltered (n=1882)
Altered (n=30)
Breast Tumor
B
PR Status
% of breast tumor samples
100
80
60
40
20
0
Unknown
Negative
Positive
p=3.314e-3
q=0.0199
Unaltered (n=1885)
Altered (n=33)
Breast Tumor

**Figure S3.**

A

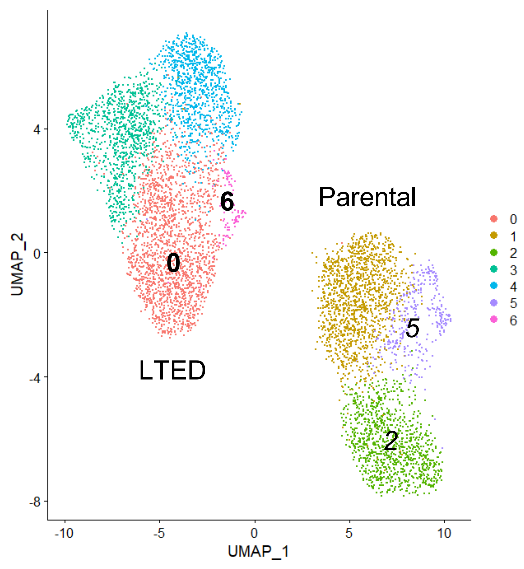


B

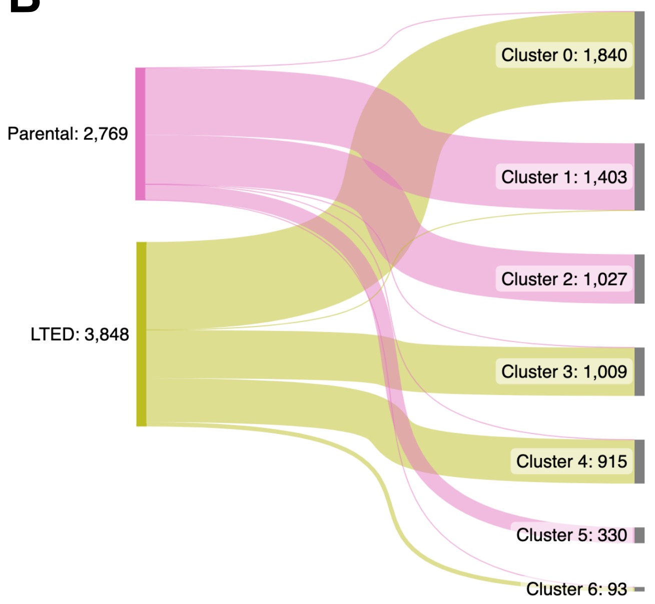


C

| | **Cluster 0** | Cluster 1 | *Cluster 2* | Cluster 3 | Cluster 4 | *Cluster 5* | **Cluster 6** |
|---|---|---|---|---|---|---|---|
| Parental | **6** | 1399 | *1027* | 2 | 1 | *330* | **4** |
| LTED | **1834** | 4 | *0* | 1007 | 914 | *0* | **89** |
| Total Cells | **1840** | 1403 | *1027* | 1009 | 915 | *330* | **93** |

D

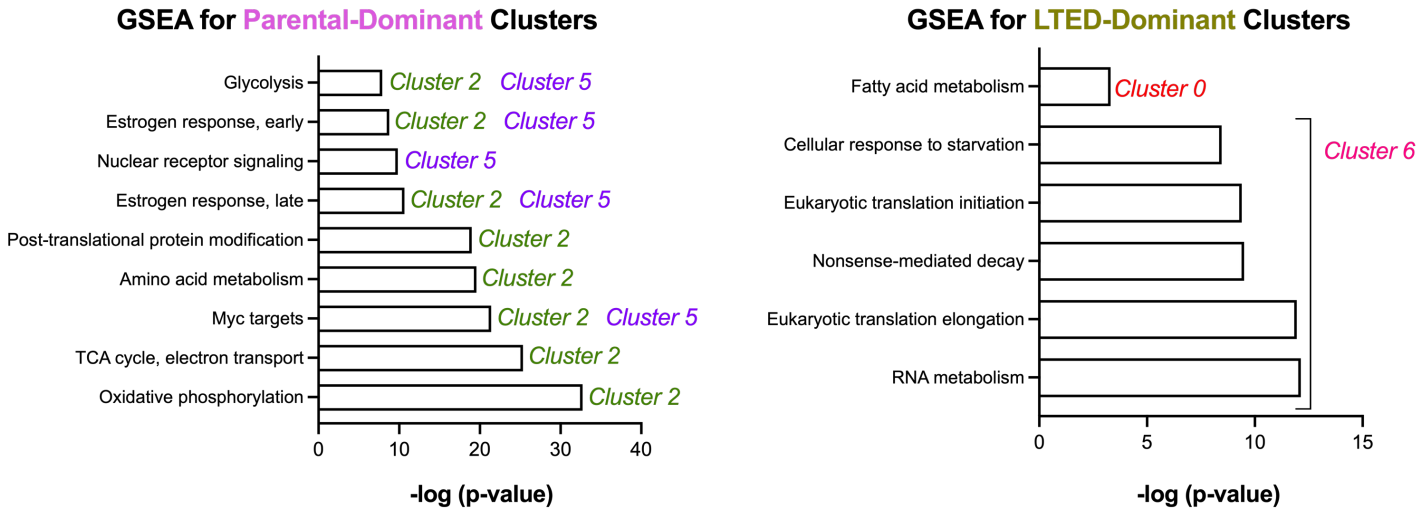

### Figure S4.

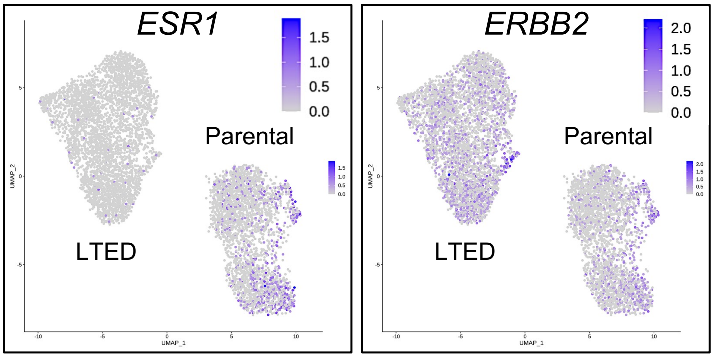

ESR1
1.5
1.0
0.5
0.0
Parental
LTED
UMAP_2
UMAP_1
ERBB2
2.0
1.5
1.0
0.5
0.0
Parental
LTED
UMAP_2
UMAP_1

**Figure S5.**

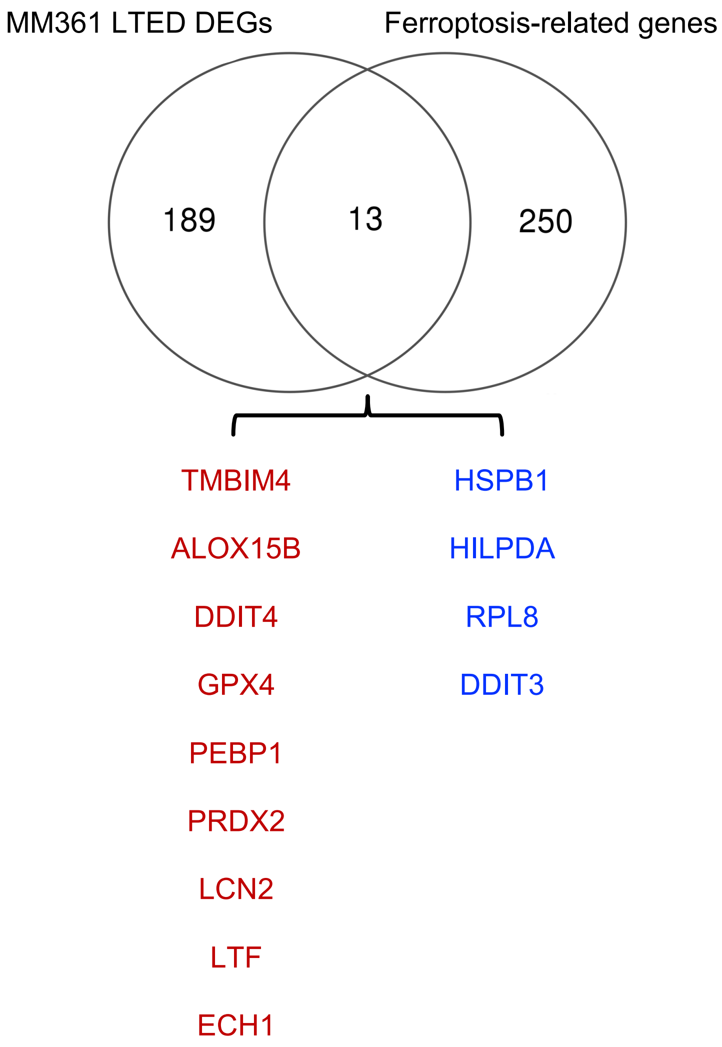
MM361 LTED DEGs
Ferroptosis-related genes
189
13
250
TMBIM4
ALOX15B
DDIT4
GPX4
PEBP1
PRDX2
LCN2
LTF
ECH1
HSPB1
HILPDA
RPL8
DDIT3

**Fig S6.**

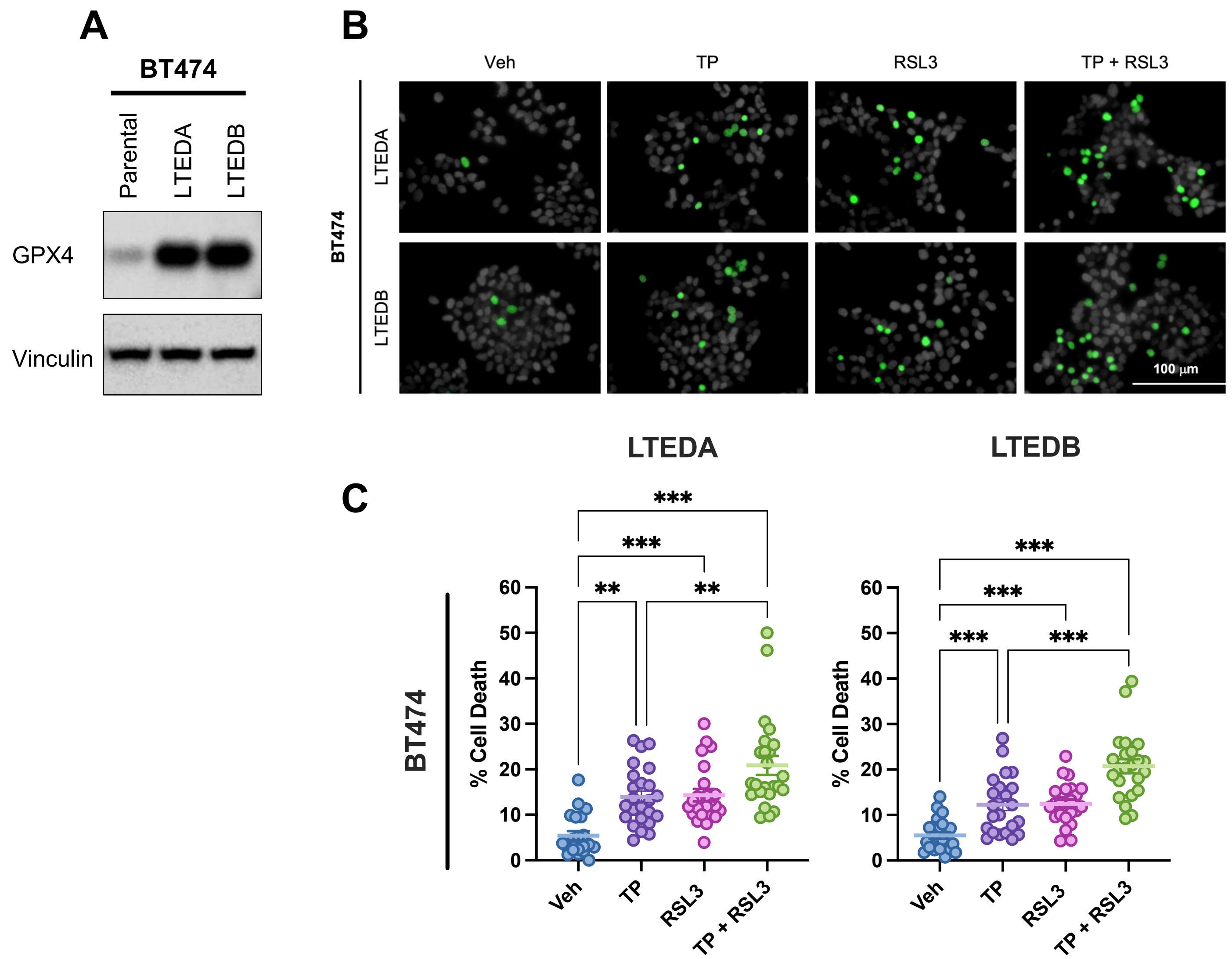

A
BT474
Parental
LTEDA
LTEDB
GPX4
Vinculin
B
Veh
TP
RSL3
TP + RSL3
BT474
LTEDA
LTEDB
100 µm
C
LTEDA
LTEDB
BT474
% Cell Death
0
10
20
30
40
50
60
***
**
Veh
TP
RSL3
TP + RSL3

#### Figure S7.

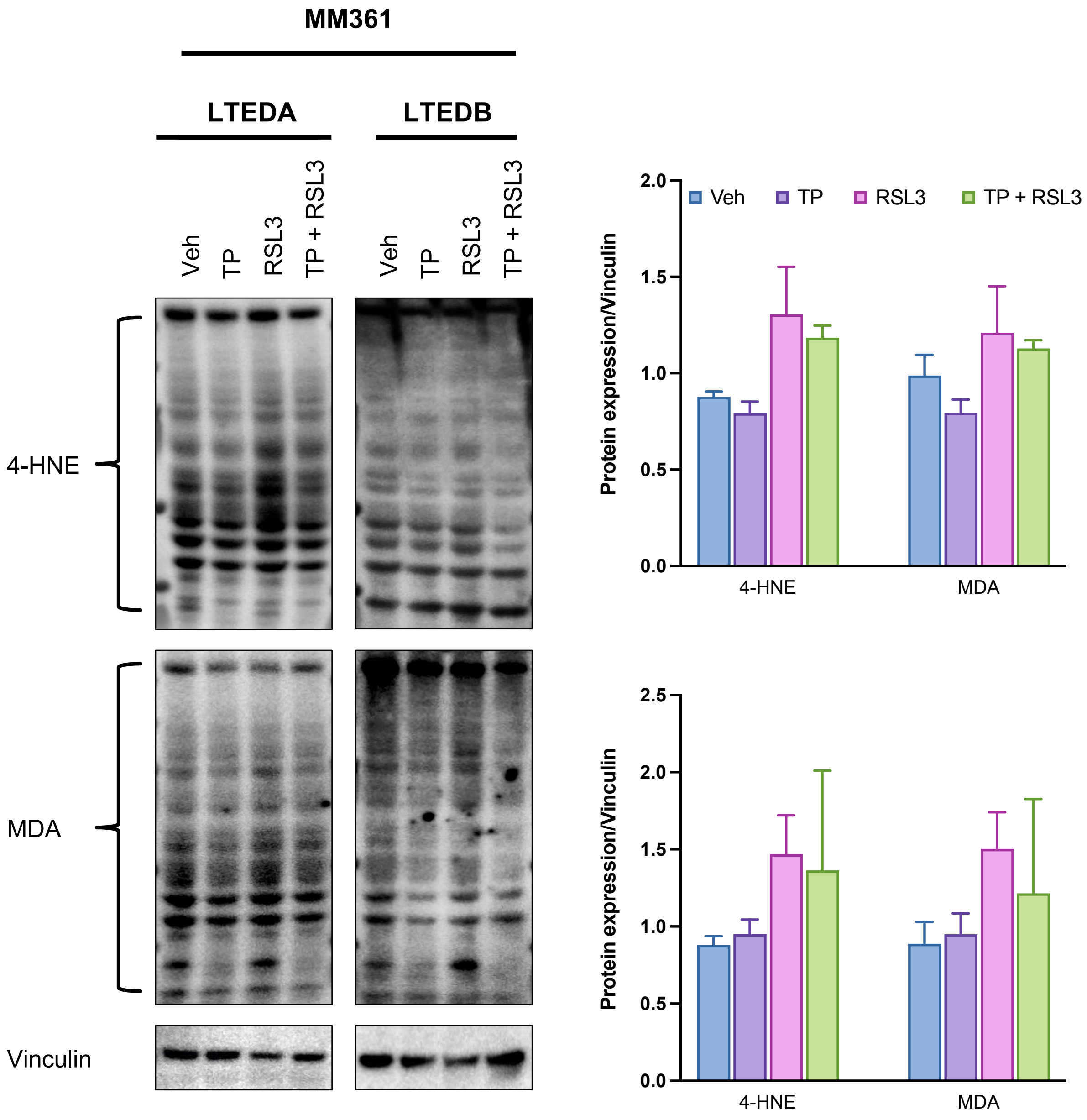

MM361
LTEDA
LTEDB
Veh
TP
RSL3
TP + RSL3
4-HNE
MDA
Vinculin
Protein expression/Vinculin
Veh
TP
RSL3
TP + RSL3
2.0
1.5
1.0
0.5
0.0
4-HNE
MDA
2.5
